# The leptin receptor has no role in delta-cell control of beta-cell function in the mouse

**DOI:** 10.1101/2023.06.30.547271

**Authors:** Jia Zhang, Kay Katada, Elham Mosleh, Andrew Yuhas, Guihong Peng, Maria L. Golson

## Abstract

Leptin inhibits insulin secretion from isolated islets from multiple species, but the cell type that mediates this process remains elusive. Mouse models have been used to explore this question. Ablation of the leptin receptor (Lepr) throughout the pancreatic epithelium results in altered glucose homeostasis, ex vivo insulin secretion, and calcium dynamics. However, the removal of Lepr from neither alpha nor beta cells mimics these results. Because Lepr is enriched in the delta cells of human islets, we used a mouse model to test whether delta cells mediate the diminished glucose-stimulated insulin secretion in response to leptin. However, ablation of Lepr within mouse delta cells had no impact on glucose homeostasis or insulin secretion. We further demonstrate that Lepr is not appreciably expressed within mouse delta cells.

## Introduction

Leptin, a hormone secreted by fat, limits hunger and promotes physical activity by activating its cognate receptor (1). It exerts these effects primarily through the hypothalamus. The leptin receptor (LEPR) is also expressed in immune cells, pericytes, and endothelial cells and affects various immune processes and vessel constriction (2,3). In addition to these roles, multiple labs have reported that leptin directly inhibits insulin secretion from isolated mouse, human, and rat islets (4-8). Moreover, deletion of the leptin receptor (Lepr) throughout the pancreatic epithelium using Pdx1-Cre alters glucose homeostasis (9). Pdx1-Cre;Leprfl/fl mice fed a normal chow diet display improved glucose tolerance, while those fed a high-fat diet display glucose intolerance (9). Considering that leptin inhibits insulin secretion, deleting Lepr would presumably promote insulin secretion, as observed on a normal chow diet. Reconciling the data on the high-fat diet is more difficult, but perhaps tonically increased insulin secretion in times of high metabolic demand leads to beta-cell exhaustion and thus the observed glucose tolerance.

While the altered glucose homeostasis in Pdx1-Cre;Leprfl/f suggests that the cell responsible for this phenotype has an epithelial origin, narrowing down the exact cell type has proven difficult. Two different models of Lepr deletion within beta cells have yielded conflicting results. Rat insulin promoter (RIPMgn)-Cre;Lepr mice display hyperglycemia and reduced insulin secretion, but they also have high adiposity due to RIP-CreMgn activity in the brain (10,11); therefore, the disrupted glucose homeostasis is likely a secondary effect to obesity associated with loss of Lepr in the hypothalamus. A second model of beta-cell Lepr ablation using the Insulin 1 (Ins1-Cre) driver resulted in very little change in blood glucose regulation, and only in females (12). In addition, alpha-cell ablation of Lepr using either Glucagon (Gcg)Cre or Gcg-CreER causes no change in glycemia (12,13).

Another primary islet endocrine cell type is the somatostatin-secreting delta cell. Somatostatin is an autocrine/paracrine hormone that inhibits secretion from adjacent alpha and beta cells. Single-cell RNA sequencing (scRNAseq) of human islets revealed that LEPR is highly enriched within delta cells (14). Therefore, we hypothesized that leptin acts by activating delta cells, promoting somatostatin release, thereby indirectly inhibiting beta cells (Figure 1A). We tested this concept in a mouse model of delta-cell-specific Lepr ablation.

**Fig. 1.**
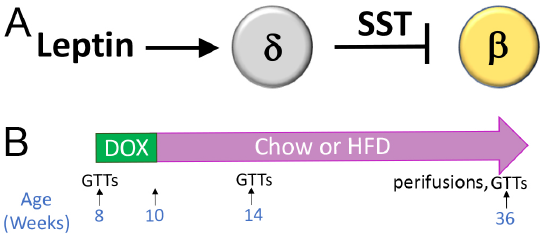
Testing whether leptin mediates inhibition of insulin secretion through the delta cell. A). In human islets, the leptin receptor is expressed almost exclusively in delta cells. We hypothesized that leptin stimulates somatostatin secretion from delta cells, thereby inhibiting insulin secretion through the activation of somatostatin receptors in beta cells. B) Mouse model: Sst(rtTA/+);Tet-O-Cre;Leprfl/fl and littermate control mice were placed on doxycycline from eight weeks to ten weeks of age. Mice were assigned to a high-fat or normal chow diet at ten weeks of age. GTTs were performed at eight, 14, and 36 weeks of age (four weeks or six months after being assigned to chow or high-fat diet).

## Methods

### Mice

Sst-rtTA (15), Tet-O-Cre (16), and Leprfl/fl (17) mice and genotyping have been described previously. 2 percent doxycycline in water was administered at eight weeks of age for two weeks. Mice were fed chow or 60 percent HFD (Research Diets) ad libitum and maintained on a 12-hour light/dark cycle. Control mice were littermates of mutants and all had one allele of Sst(rtTA) and included Sst(rtTA/+);Tet-O-Cre;Lepr+/fl and SstrtTA;Leprfl/fl mice. All procedures followed the Institutional Animal Care and Use Committee guidelines at the University of Pennsylvania or Johns Hopkins University.

### Glucose Tolerance Tests, Mouse Islet Isolations, and Perifusions

For glucose tolerance tests (GTTs), mice were fasted overnight for 16 hours, and blood glucose was measured with an AlphaTRAK glucometer. Mice were then injected intraperitoneally with 1g/kg or 2g/kg glucose in sterile 1X PBS; blood glucose was measured 15, 30, 60, 90, and 120 minutes after injection. Islets were isolated by collagen digestion followed by Ficoll separation and hand-picking, as previously described (18). Perifusions were performed by the University of Pennsylvania Pancreatic Islet Cell Biology Core.

### Recombination Analysis

Pancreata were fixed in 4 percent PFA for 4 hours, rinsed in 1X PBS, and embedded in paraffin. 4-micron sections were placed on PermaFrost slides. One to four sections were then scraped with a clean razor into a sterile microcentrifuge tube and digested overnight with 100 microliters of Proteinase K solution before standard PCR, using 5-10 microliters of heat-inactivated digested sections. Primers for recombination analysis were previously described (19).

### qPCR

Primers (Lepr F1 5’-GTT TCA CCA AAG ATG CTA TCG AC-3’; Lepr R1 5’GAG CAG TAG GAC ACA AGA GG-3’) for Lepr were tested on hypothalamus cDNA before analyzing Lepr expression in islets. RNA was isolated and cDNA was generated from whole islets or hypothalamus using Trizol plus the Qiagen RNeasy kit, as previously described (20). qPCR was performed as previously described (20).

### Bulk RNAseq

Human islets were obtained from the Integrated Islet Distribution Program (IIDP). Islet preparation and RNA extraction after intracellular marker cell sorting has been described previously (21). Briefly, islets from five non-diabetic human donors (Table 1) were suspended into a single-cell solution using 0.05 percent Trypsin, then fixed in 4 percent PFA, 0.1 percent saponin, and 1:100 RNasin (Promega). Cells were then indirectly fluorescently immunolabeled against Somatostatin and Insulin and sorted. RNA was extracted with a Recoverall Total Nucleic Acid kit (Life Technologies). RNA library was prepared and sequenced as previously described (22).

**Table 1.** Donor characteristics

| Donor | Sex | Age | BMI | Race/Ethnicity |
| --- | --- | --- | --- | --- |
| 1 | M | 35 | 23.6 | White |
| 2 | F | 53 | 20.5 | White |
| 3 | F | 36 | 35.7 | Hispanic |
| 4 | F | 24 | 35.3 | White |
| 5 | M | 25 | 24.3 | Black |

### scRNAseq

scRNAseq data was downloaded from PANCDB (pmacs.hpap.upenn.edu) on March 31, 2023, using celltype calling performed by the Human Pancreas Analysis Program (HPAP). Non-diabetic, autoantibody-negative male and female donors >17 years of age were included in our analysis. Differential expression analysis was performed as previously described (18).

## Results

Differential expression between bulk-sorted human beta and delta cells. Current sorting protocols for live human islet cells cannot reliably distinguish delta cells from beta cells (23,24). Therefore, gene expression differences have been reported between bulk-sorted human alpha and beta cells (25) but not between bulk-sorted human beta and delta cells. To separate these two closely related cell types, we fixed islets from five non-diabetic human donors, suspended them in a single-cell suspension, permeabilized them, and indirectly immunolabeled them with antibodies against Insulin and Somatostatin. These cells then underwent fluorescence-activated cell sorting. RNA was extracted from the purified beta- and delta-cell populations and subjected to RNAseq and analysis. As expected, known lineage-specific genes such as SST and HHEX were strongly enriched in delta cells (Table 2), while genes such as INS and MEG3 were more highly expressed in beta cells (Table 3).

**Table 2.** Top enriched genes in sorted delta cells compared to beta cells.

| Gene Symbol | Fold Enrichment | FDR |
| --- | --- | --- |
| SST | 283.354336 | 3.2099E-145 |
| ERBB4 | 231.194075 | 1.56518E-71 |
| SLITRK6 | 182.211784 | 4.29086E-57 |
| SALL1 | 173.469728 | 3.27556E-71 |
| AGTR1 | 126.263771 | 3.30735E-43 |
| MS4A8B | 121.274614 | 4.99604E-70 |
| HHEX | 115.78778 | 1.50192E-81 |
| KCNIP1 | 99.9023306 | 7.06592E-51 |
| CBLN4 | 98.1777133 | 4.44692E-39 |
| EPHA6 | 92.2108402 | 8.98006E-33 |
| TMEM132E | 86.6689936 | 3.49776E-31 |
| ODZ2 | 74.9405509 | 1.00547E-60 |
| IGSF10 | 71.1360064 | 2.32374E-60 |
| DCHS2 | 70.6242855 | 1.42405E-62 |
| FLRT1 | 70.407892 | 9.4954E-52 |
| DRD2 | 70.2015891 | 5.55123E-33 |
| GABRA1 | 67.364591 | 2.03333E-26 |
| PDE2A | 64.9836722 | 4.78243E-23 |
| SLC5A8 | 64.1198737 | 4.02285E-26 |
| LRFN5 | 63.8498264 | 9.00996E-94 |
| NTNG1 | 62.2271069 | 1.84028E-46 |
| FAM5C | 60.5572346 | 1.44708E-16 |
| LRRC4C | 55.5375203 | 4.22091E-18 |
| BCHE | 55.1893356 | 3.90434E-54 |
| LPHN2 | 53.0600885 | 4.78117E-82 |
| C1orf173 | 52.1434247 | 5.11576E-65 |
| KCNT2 | 50.3648355 | 8.46998E-27 |
| FHOD3 | 48.5091315 | 6.404E-65 |
| SRRM4 | 47.7582127 | 4.48161E-23 |
| PREX2 | 46.0390162 | 1.45321E-27 |
| PPFIA2 | 44.9601447 | 1.52849E-69 |
| LEPR | 44.844213 | 2.30687E-97 |
| RYR1 | 44.183812 | 3.95159E-29 |
| GABRA5 | 43.5690639 | 7.12644E-21 |
| KCNC2 | 40.9824238 | 1.66405E-27 |
| LOC100233209 | 40.6414705 | 1.11879E-21 |
| ODZ1 | 40.6015994 | 1.99714E-29 |
| SORCS1 | 39.5170708 | 8.74982E-16 |
| LEPR | 39.2396012 | 2.91084E-77 |
| PDE1C | 38.7210966 | 4.91781E-23 |
| FAM113B | 38.5093709 | 2.17406E-27 |
| ARHGEF38 | 35.7514589 | 1.99714E-29 |
| CPA4 | 35.7382946 | 5.60967E-32 |
| SEMA3E | 35.5044204 | 4.55075E-49 |
| MLPH | 35.392378 | 6.38692E-43 |
| SERPINA1 | 35.0834445 | 3.17001E-26 |
| RGS6 | 34.7928073 | 1.3526E-28 |
| GABRG2 | 34.2872608 | 2.23566E-10 |
| LAMA1 | 32.8074723 | 3.07579E-19 |

**Table 3.** Top enriched genes in sorted beta cells compared to delta cells.

| Gene Symbol | Fold Enrichment | FDR |
| --- | --- | --- |
| PPM1E | 77.3401176 | 1.44869E-38 |
| PCA3 | 60.7783158 | 6.30411E-32 |
| ZNF114 | 56.2000104 | 2.03476E-27 |
| SNORD113-1 | 56.1416837 | 2.3718E-26 |
| PRKCH | 54.2072419 | 4.53726E-36 |
| ZPLD1 | 53.8064123 | 6.9859E-25 |
| TRIM9 | 51.9466875 | 2.03706E-29 |
| TSPAN1 | 49.7902167 | 1.58607E-52 |
| MEG8 | 44.8834693 | 1.5268E-26 |
| MEG3 | 41.8423225 | 1.90814E-21 |
| IGSF11 | 41.7490255 | 8.9035E-13 |
| WSCD2 | 38.6772911 | 6.7188E-50 |
| IGFBP3 | 37.994922 | 1.71278E-19 |
| SNORD114-31 | 37.0304821 | 1.25214E-48 |
| TDRG1 | 35.9852021 | 1.89274E-66 |
| TMEM108 | 35.5838299 | 4.82458E-41 |
| TFF3 | 35.1481675 | 4.49843E-92 |
| LOC285501 | 34.4432135 | 5.43917E-15 |
| LY75 | 31.8524874 | 3.51209E-24 |
| OTOGL | 31.2755596 | 4.12386E-13 |
| PTGS2 | 30.9494755 | 6.76815E-28 |
| TNFRSF11A | 30.8657815 | 2.33819E-50 |
| ROBO2 | 29.024644 | 1.3592E-23 |
| LPAR4 | 28.8794137 | 1.96509E-26 |
| SLC2A2 | 27.9735477 | 2.91653E-20 |
| PPP2R2C | 27.0457367 | 2.13994E-19 |
| HOPX | 26.8617054 | 1.45504E-17 |
| ZNF385D | 26.5909198 | 2.98103E-21 |
| SNORD114-12 | 24.9059593 | 2.65463E-36 |
| NCAN | 23.9614853 | 1.52079E-20 |
| MFI2 | 23.8734734 | 1.13795E-21 |
| TCERG1L | 23.8421319 | 1.22641E-20 |
| SNORD114-11 | 23.230222 | 1.79034E-27 |
| SNORD114-7 | 23.2297229 | 8.24218E-45 |
| KCNG3 | 22.0542854 | 4.43018E-08 |
| SLC6A6 | 21.9826146 | 8.06183E-43 |
| LRFN2 | 21.4123197 | 1.34912E-15 |
| FOXQ1 | 21.250562 | 4.17158E-08 |
| SNORD114-2 | 20.5709127 | 2.10688E-05 |
| HAPLN4 | 20.5445129 | 1.13442E-11 |
| INS-IGF2 | 20.3647453 | 9.97745E-10 |
| SPAG6 | 19.6697994 | 1.26075E-30 |
| RPH3A | 19.5420834 | 4.53605E-10 |
| TM6SF2 | 19.5391075 | 1.06378E-26 |
| LOC145837 | 19.5333186 | 4.58414E-11 |
| HS6ST2 | 19.2960624 | 1.51035E-09 |
| RGS16 | 19.0761522 | 4.93904E-27 |
| TNNI1 | 19.0404075 | 1.84728E-37 |
| MAF | 18.7190384 | 1.24332E-14 |
| HHATL | 18.698355 | 5.98751E-08 |
| INS | 18.6035554 | 1.01903E-09 |

An additional gene enriched within delta cells was LEPR, in agreement with a previous scRNAseq study (14). LEPR appears twice, as both a full-length (NM002303, >39-fold enriched) and a truncated (NM001198689, >44-fold enriched) isotype. Since our bulk-sorted RNA sequencing was performed on a mixture of male and female donors, we also examined gene expression differences between male and female beta and delta cells using publicly available scRNAseq data from the Human Islet Research Network’s Human Pancreas Analysis Program (Tables 4-5; (26)). The genes enriched within male and female delta cells compared to beta cells were similar, although the magnitude of differential expression varied. LEPR transcripts were more abundant in delta cells compared to beta cells by 1.49-fold and 1.33-fold with an adjusted p-value of 3.80 × 10-308 or 9.99 x10-311 in females and males, respectively. Although LEPR was not among the top 50 enriched delta-cell genes in males, it was in females (Tables 4-5).

**Table 4.**
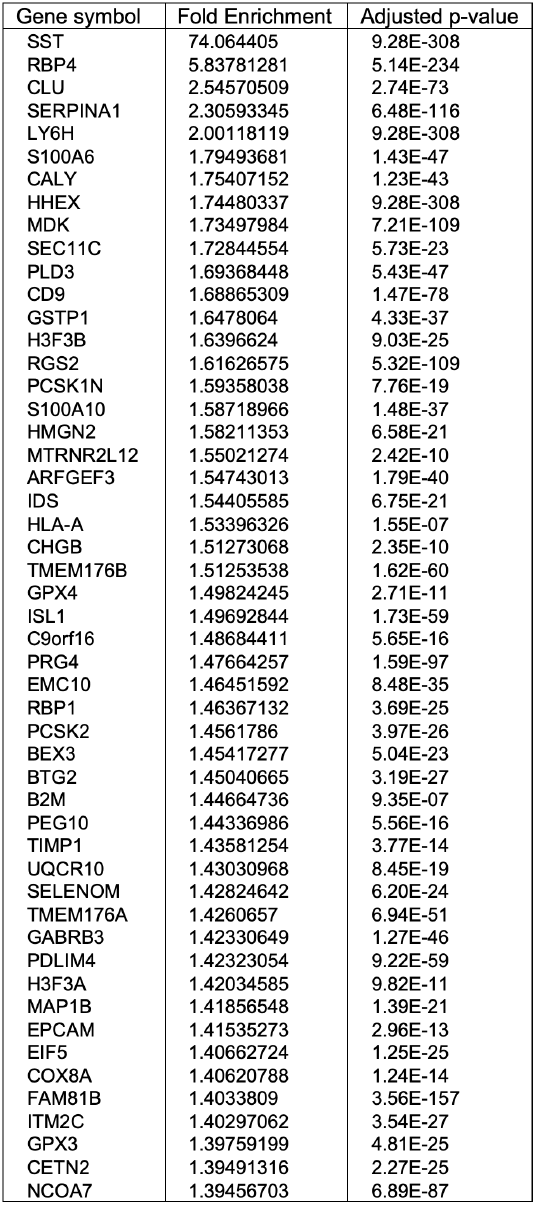
Top delta-cell enriched genes compared to beta cells from male HPAP

**Table 5.**
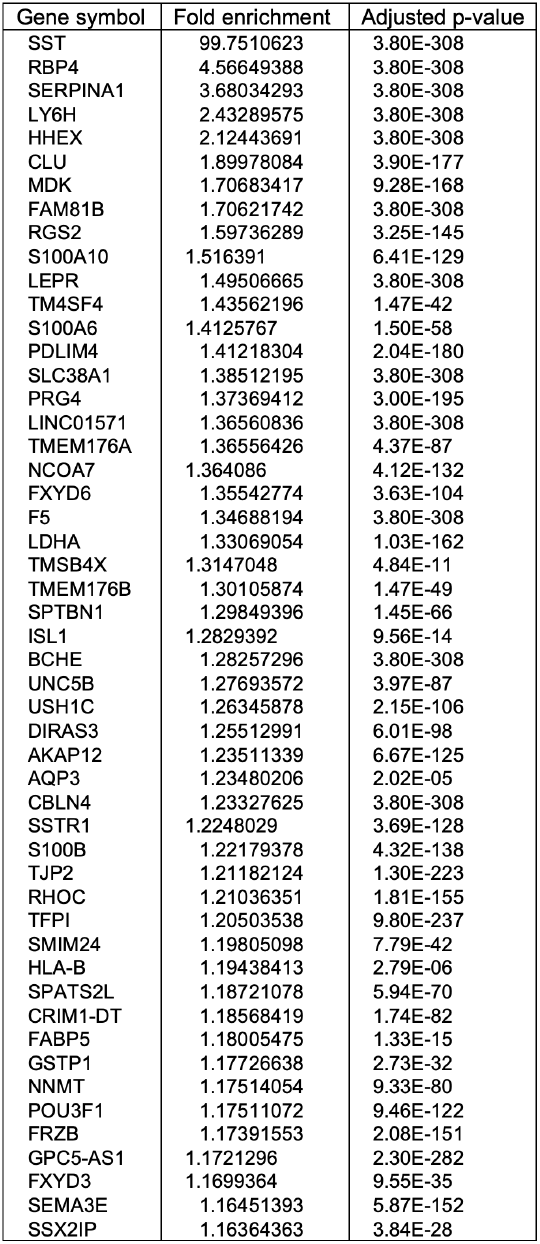
Top delta-cell enriched genes compared to beta cells from female HPAP scRNAseq samples.

Delta-cell-specific Lepr ablation in the mouse results in normal glucose homeostasis and insulin secretion. The fact that ablation of the Lepr within the mouse pancreatic epithelium results in alterations in glucose homeostasis, whereas ablation within alpha or beta cells has no effect, combined with the enrichment of LEPR in human delta cells, suggested that delta cells might mediate the effects on insulin secretion observed in response to leptin treatment. Therefore, we derived a mouse model (SST(rtTA/+);Tet-O-Cre;Leprfl/fl mice) to test this hypothesis (Figure 1B). Mice were treated with doxycycline for two weeks, followed by a four-week washout to exclude any metabolic side effects from doxycycline treatment. In addition, since doxycycline would induce recombination of Lepr in any cells expressing somatostatin, we also Weight was tracked weekly. No difference in weight was observed between SST(rtTA/+);Tet-O-Cre;Leprfl/fl and control mice on high-fat or chow diet (not shown). Histological sections were immunolabeled with insulin, glucagon, and somatostatin, and no differences in beta-cell mass, endocrine cell ratios, or islet morphology were noted (data not shown). Analysis of Lepr gene recombination and RNA expression. We next examined Lepr locus recombination using DNA extracted from pancreatic sections from 36-weekold Sst(rtTA/+);Tet-O-Cre;Lepr+/fl, Sst(rtTA/+);Tet-OCre;Leprfl/fl, Ss t(rtTA/+);Lepr+/fl, and Sst(rtTA/+);Leprfl/fl mice that had been treated with doxycycline from eight to ten weeks of age. The Leprfl allele was designed using loxP elements surrounding the first exon of Lepr (17). We used primers that flanked either one loxP element or both loxP elements, including the first exon of Lepr (Figure 2A). Primers flanking one loxP element amplify unrecombined DNA, as would be expected from non-delta cells. Amplification with these primers cells resulted in one (Leprfl/fl mice) or two bands (Lepr+/fl) in all samples (Figure 2B), with one band indicating homozygosity of the floxed allele and two bands indicating one wild-type and one floxed allele. The presence of the bands of the unrecombined allele in Cre+ animals reflects the large number of non-delta cells in the endocrine pancreas.

**Fig. 2.**
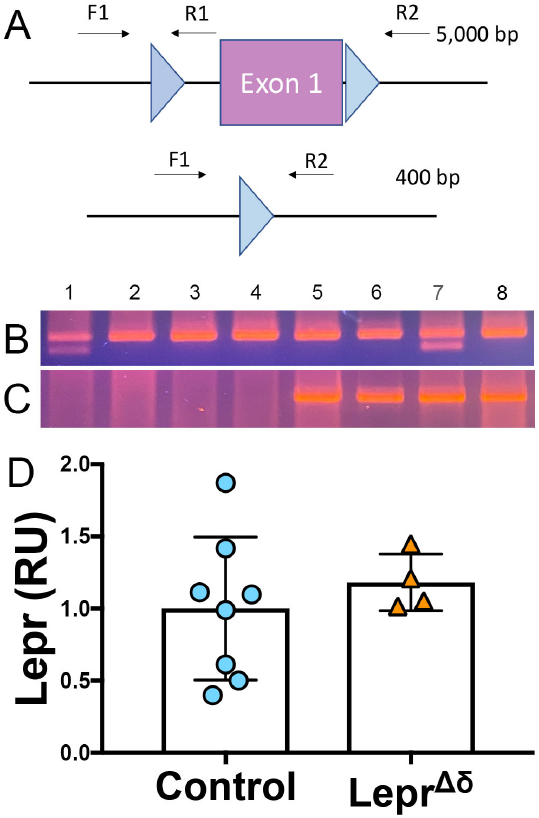
Lepr recombination and expression analysis. A) Schematic of Leprfl allele and primers used to assess DNA recombination of the loxP elements. B-C) PCR was performed on DNA collected from fixed p ancreatic s ections. F 1/R1 primers amplify unrecombined DNA (B), while F1/R2 primers result in a 400bp product after Leprfl recombination (C). All samples tested had one allele of SstrtTA. Samples 1 and 7 were heterozygotes for Leprfl. Samples 1-4 were Tet-O-Cre-negative, while Samples 5-8 were Tet-O-Cre-positive. D) qPCR analysis of Lepr expression in whole islets from Leprfl or control mice.

In the absence of recombination, primers flanking both loxP elements span an 5kb distance, which is too long for amplification using standard PCR reagents (Figure 2A). A recombined Lepr gene results in an approximately 400-bp amplicon. In the absence of Tet-O-Cre, no band was observed. All samples from mice harboring at least one allele of Sst-rtTA, Tet-O-Cre, and Leprfl yielded the 400-bp band expected when recombination occurs (Figure 2C), reflecting recombination of the loxP allele in delta cells. To test whether Lepr expression was reduced in Sst-rtTA/+;Tet-O-Cre;Leprfl/fl delta cells, whole islets were isolated from 14-week-old Sst(rtTA/+);Tet-O-Cre;Leprfl/fl and control mice for quantitative RT-PCR analysis. No difference was observed in Lepr expression between Sst(rtTA/+);Tet-O-Cre;Leprfl/fl and control mice (Figure 2D). Others have reported decreased Lepr expression when this Leprfl mouse was used for deletion in other tissues (17); however, given that delta cells make up less than 5 percent of endocrine cells in mice, if other islet cell types also express appreciable amounts of Lepr mRNA, detecting deletion would be difficult.

Glucose tolerance is unaltered in SST(rtTA/+);Tet-OCre;Leprfl/fl mice. Intraperitoneal GTTs were performed with 2g/kg glucose at 8 weeks, 14 weeks, and 36 weeks of age, corresponding to pre-doxycycline, and then four weeks or six months after doxycycline removal. No difference was observed in glucose tolerance in male (Figure 3) or female (Figure 4) SST(rtTA/+);Tet-O-Cre;Leprfl/fl mice mice at any time point, whether the mice were on a standard chow or high-fat diet. To rule out the possibility that we were missing subtle differences, GTTs were also performed on males and females on a HFD diet using 1g/kg glucose (Figure 5). Again, no alterations in glucose tolerance were observed. Insulin secretion was examined by islet perifusion at 36 weeks of age. Islets from male and female SST(rtTA/+);Tet-O-Cre;Leprfl/fl mice mice displayed no significant differences in insulin secretion compared to controls (Figure 6A-D).

**Fig. 3.**
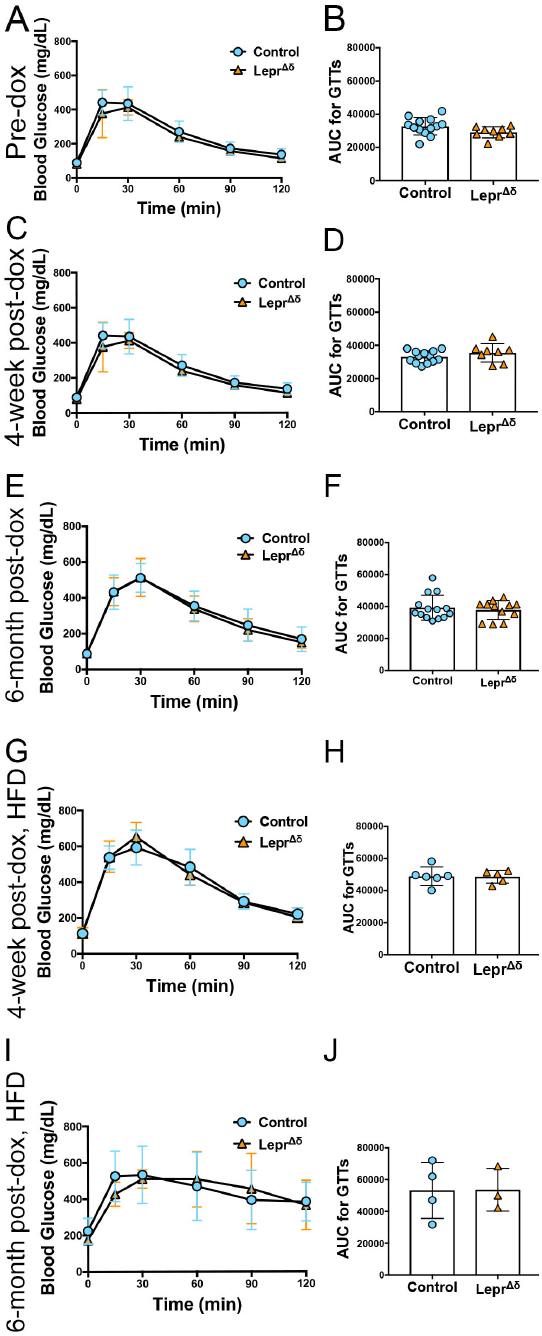
Sst(rtTA/+);Tet-O-Cre;Leprfl/fl male mice display normal glucose tolerance on chow and high-fat diet. A-J) GTTs (A, C, E, G, I) and corresponding area-underthe-curve analysis (B, D, F, H, J) for male SST(rtTA/+);Tet-O-Cre;Leprfl/fl mice prior to doxycycline administration (A-B), at 14 weeks of age and four weeks after doxycycline removal and placement on a chow diet (C-D), at 36 weeks and six months after doxycycline removal and placement on a chow diet (E-F), at 14 weeks of age and four weeks after doxycycline removal and placement on a high-fat diet (G-H), and at 36 weeks of age and six months after doxycycline removal and placement on a high-fat diet.

**Fig. 4.**
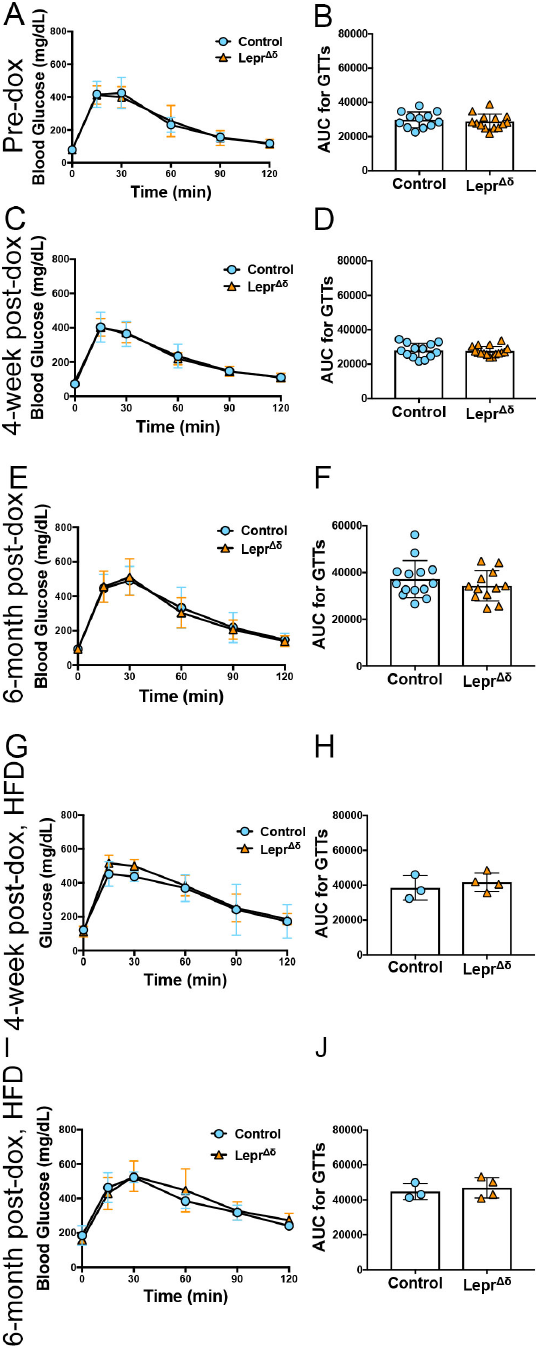
SST(rtTA/+);Tet-O-Cre;Leprfl/fl mice female mice display normal glucose tolerance on chow and high-fat diet. A-J) GTTs (A, C, E, G, I) and corresponding areaunder-the-curve analysis (B, D, F, H, J) for female SST(rtTA/+);Tet-O-Cre;Leprfl/fl mice prior to doxycycline administration (A-B), at 14 weeks of age and four weeks after doxycycline removal and placement on a chow diet (C-D), at 36 weeks and six months after doxycycline removal and placement on a chow diet (E-F), at 14 weeks of age and four weeks after doxycycline removal and placement on a highfat diet (G-H), and at 36 weeks of age and six months after doxycycline removal and placement on a high-fat diet.

**Fig. 5.**
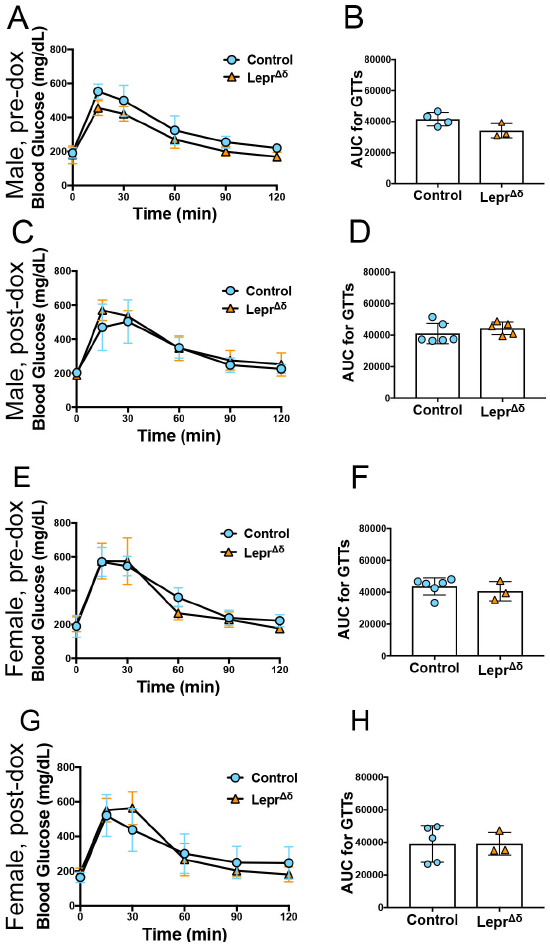
Male and female SST(rtTA/+);Tet-O-Cre;Leprfl/fl mice on a high-fat diet display normal glucose tolerance in response to a reduced glucose load. Glucose tolerance tests using 1g/kg glucose were performed before doxycycline treatment (A-B, E-F) and at 14 weeks of age in male (C-D) and female (E-H) SST(rtTA/+);TetO-Cre;Leprfl/fl and control mice on a HFD diet starting at 10 weeks of age. Glucose curves (A, C, E, and G) and area-under-the-curve (B, D, F, and H) are shown.

**Fig. 6.**
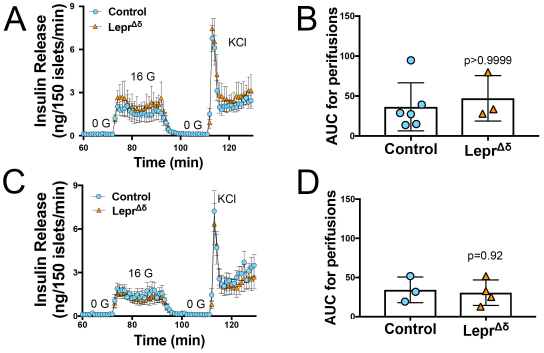
Islets from SST(rtTA/+);Tet-O-Cre;Leprfl/flmice display normal insulin secretion in response to high glucose. Isolated islets from 36-week old male (A-B) and female (C-D) SST(rtTA/+);Tet-O-Cre;Leprfl/fl and control mice were subjected to perifusions (A,C) at 0 mM glucose (0 G), 16 mM glucose (16 G), and with KCl. Both males (A-B) and females (C-D) were examined. Area-under-the-curve for 16 mM glucose is shown (B,D).

Lepr is highly expressed in endothelial cells, but not delta cells, within the mouse islet. We, therefore, examined Lepr expression in mouse alpha, beta, and delta cells using a published dataset built from fluorescently sorted populations of live lineage-traced cells. Lepr was not detected in mouse alpha, beta, or delta cells (Figure 7A, (27)). This result starkly contrasts the transcription factor Pdx1, which is detectable at the RNA level in each population (Figure 7B) despite the limitation of PDX1 protein expression in the mature pancreas to beta and delta cells.

**Fig. 7.**
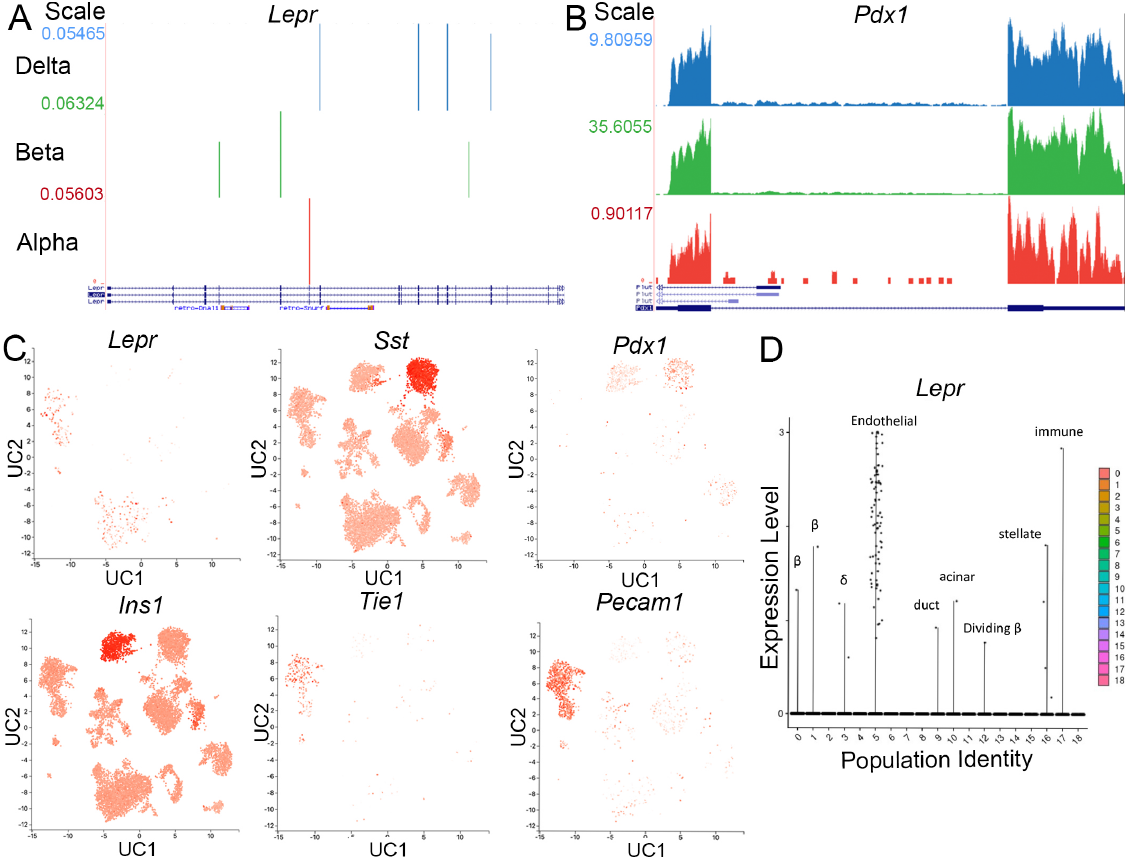
Lepr is not expressed in mouse delta cells. A-B) RNAseq reads for Lepr (A) and Pdx1 (B) in sorted alpha, beta, and delta cells were obtained from publicly available data (27). C) scRNAseq expression in mouse islets cells for Lepr, Sst, the ductal, beta-and delta-cell marker Pdx1, Ins1, and two endothelial markers, Tie1 and Pecam, was obtained from the public database PanglaoDB (https://panglaodb.se, SRA745567). UC: UMAP component. D) Expression of Lepr in identified cell populations from a second published mouse islet scRNAseq dataset (18).

Examination of a publicly available scRNAseq dataset confirmed a lack of Lepr expression within the mouse islet endocrine population (Figure 7C). Most Lepr-positive cells were observed in endothelial cell clusters (Figure 7C). Our scRNAseq data (18) also demonstrated that most Lepr-positive cells express endothelial cell markers (Figure 7D). In addition, Lepr expression was detected in only 0.01 percent of islet endocrine cells.

## Conclusions

Multiple labs have demonstrated that leptin decreases insulin secretion from isolated islets. In addition, beta cells from control mice hyperpolarize in response to leptin, but this response is lost in beta cells from global Lepr null mice (28).

These data strongly indicate that a cell type residing within the islet mediates altered insulin secretion in response to leptin. However, pinpointing this cell type has proven to be challenging.

Pdx1-Cre is expressed throughout the pancreatic epithelium; this Cre would induce recombination of Lepr in the islet endocrine and acinar cells but not in the immune, endothelial, or smooth muscle cells that might be isolated along with the islet (29). Pdx1-Cre;Leprfl/fl male and female mice on a normal chow diet exhibited improved glucose tolerance, normal insulin tolerance, and increased fasting serum insulin (9). Isolated Pdx1-Cre;Leprfl/fl islets display increased Ca2+ influx and insulin secretion compared to Leprfl/fl controls without Cre. Furthermore, leptin treatment represses insulin secretion in isolated islets from control but not Pdx1-Cre;Leprfl/fl mice (9).

These data suggest that pancreatic epithelial cells are responsible for the depression of glucose-stimulated insulin secretion in response to leptin in isolated islets. However, ablation of Lepr in alpha cells using an inducible Glucagon-Cre (12), in beta cells using Ins1-Cre (12), or in delta cells (demonstrated here) does not result in altered glucose tolerance or insulin secretion. Additionally, bulk RNAseq of sorted mouse alpha, beta, or delta cells cannot detect Lepr (27). Furthermore, in this manuscript, we show by scRNAseq that Lepr can be detected in only an exceedingly small percentage of endocrine or acinar cells in mice. Genes with low expression are sometimes detectable in only a small percentage of cells using scRNAseq. One such gene is Pdx1, which can be observed in both beta- and delta-cell clusters, but in many fewer cells than somatostatin or insulin (Figure 7C). Despite these drop-outs in scRNAseq, Pdx1 is readily detectable in sorted beta cells (Figure 7B). In addition, in a Lepr-Cre;TdTomato reporter mouse, no fluorescence w as d etected i n i slet endocrine cells (30). Together, these data suggest that Lepr is largely or wholely absent from mouse islet endocrine cells.

It has been reported previously that LEPR is expressed in mouse T cells (31,32), endothelial cells (33), and smooth muscle cells (33). Our scRNAseq also demonstrates Lepr expression within islet endothelial cells. However, mice with ablated Lepr within endothelial cells display unchanged ad libitum-fed blood glucose (19). Unfortunately, a more systematic analysis of glucose homeostasis was not performed in these mice.

Endothelial cells regulate beta-cell function and insulin release through factors such as Endothelin-1 and Thrombospondin-1 (34,35). Thus, signaling changes between endothelial and beta cells in response to Leptin treatment may mediate decreased insulin secretion in mouse islets. However, Pdx1-Cre should not cause recombination within endothelial cells; thus, deletion within the endothelial cells is unlikely to underlie the altered insulin secretion in the Pdx1-Cre;Leprfl/fl mo del. Moreover, in human islets, LEPR is detectable in scant endothelial cells by scRNAseq (data not shown); thus, in human islets, any difference in insulin release due to leptin is unlikely to rely on endothelial cells.

Others have suggested that the Pdx1-Cre;Leprfl/fl phenotype results from neuronal Lepr ablation since Pdx1-Cre is active in some brain cells (13). Limited brain Lepr ablation could account for the differences in glucose tolerance observed. However, this model of neuronal deletion likely does not explain the alterations in insulin secretion from and Ca2+ influx into Pdx1-Cre;Leprfl/fl isolated islets, unless long-term changes in islet function result from neuronal input. Pdx1Cre is also active in duodenal enterocytes, where Lepr is expressed, and these cells might also contribute to the altered glucose tolerance in Pdx1-Cre;Leprfl/fl mice (36,37).

While we have not settled the mystery of which cell type mediates reduced insulin secretion resulting from leptin treatment, we provide strong evidence that, at least in the mouse islet, it is not the delta cell. Further work will be necessary to determine the identity of this leptin-responsive cell in the mouse islet.

## ACKNOWLEDGEMENTS

Human pancreatic islets were provided by the NIDDK-funded Integrated Islet Distribution Program (IIDP) at City of Hope (U24-DK-098085). Human islet scRNAseq data was obtained from the NIDDK funded Human Pancreas Analysis Program (HPAP) Database, a Human Islet Research Network (HIRN) consortium (UC4-DK-112217, U01-DK-123594, UC4-DK-112232, and U01-DK-123716). This work was supported by R01-DK-110183 to MLG and a sub-award to MLG from NIH P30-DK-109525 to MAL.

